# Nonspecific steric hindrance of protein particles by lamina-associated domains

**DOI:** 10.64898/2026.05.13.724802

**Authors:** Naz Bardakcı, Aykut Erbaş, Ozan S. Sarıyer

## Abstract

Genomic organization within the nucleus is crucial for gene regulation and cell health, as disruptions in this organization are linked to genetic disorders and cancers. Recent studies suggest that molecular-scale organization of chromatin near the nuclear periphery (lamina-associated domains, LADs) affects gene regulation, providing transciptional supression, but the biophysical mechanisms of supression behind remain unclear. LADs are large heterochromatic regions near the nuclear lamina, where transcriptional factors and RNA polymerase are scarce, implying a nonspecific filtering property. Here, we investigate the steric filtering capabilities of LADs by performing coarse-grained polymer simulations. Our results show that LAD thickness can be affected by the interaction between chromatin and nuclear periphery as well as chromatin self-compaction. Regardless, the LAD layer acts as a size-selective steric partitioning environment for protein particles limiting their access to nuclear periphery. Notably, increasing bulk protein levels enhances protein access linearly. These results align with experimental observations and suggest that LADs could control the presence of transcription machinery on the periphery of the nucleus, providing a polymer-physical mechanism for gene regulation in nuclei.

## I. INTRODUCTION

In eukaryotic cells, genomic DNA is packaged into chromatin, allowing long DNA molecules to be compacted inside the micrometer-sized nucleus while remaining accessible for transcription, replication, and repair [1, 2]. Chromatin is organized in a highly nonrandom three-dimensional manner, and this spatial organization is closely linked to gene regulation [2–5]. At the molecular level, chromatin structure is controlled by nucleosome organization and histone modifications, giving rise broadly to euchromatin and heterochromatin, which differ in compaction and transcriptional activity [6–10]. In interphase nuclei, euchromatin is found more in the nuclear interior, whereas heterochromatin is preferentially enriched at the nuclear periphery [3, 4, 11].

The nuclear periphery is defined by the nuclear envelope and the underlying nuclear lamina, a protein meshwork composed primarily of Aand B-type lamins and associated membrane proteins [12–15]. Large genomic regions known as lamina-associated domains (LADs) frequently contact the lamina and constitute a major fraction of peripheral heterochromatin [11, 16, 17]. LADs typically span hundreds of kilobases to several megabases, are enriched in repressive chromatin features, and are generally associated with low transcriptional activity [11, 17–19] but can accommodate active genes as well [20– Because lamina–chromatin organization is strongly connected to genome regulation and has been implicated in developmental abnormalities, laminopathies and cancers, understanding the physical role of LADs is an important problem [16, 23].

Despite extensive genomic and epigenetic characterization of LADs, the physical basis of their repressive character remains incompletely understood. In fact, while LADs can exhibit mostly transcriptional inactivity, some genes can “escape” suppression and allow transcriptional activity, which requires interactions between DNA and relatively large protein machinery (e.g., transcription factors, RNA Polymerase) [24, 25]. Most existing explanations emphasize specific biochemical interactions between chromatin, lamins, and associated proteins [26]. However, an additional generic mechanism may also operate: the dense chromatin environment near the lamina may sterically limit the penetration of diffusing regulatory proteins. Recent live-cell and super-resolution imaging studies indicate that both euchromatin and heterochromatin can remain highly accessible in living human cells, suggesting that regulation may depend not simply on open versus closed chromatin, but also on the size and local concentration of regulatory proteins [27]. This raises the possibility of LADs acting as *size-selective steric filters*, allowing vital protein–DNA interactions at the border between LADs and the rest of the genome but reducing the local availability of larger proteins near the nuclear periphery.

In this study, we test this mechanism via coarse-grained molecular dynamics simulations of chromatin near the nuclear lamina. We model LADs as polymeric euchromatin and heterochromatin segments adsorbed onto a lamina-like surface and regulatory proteins as freely diffusing, spherical particles with tunable size. By varying chromatin–lamina affinity, chromatin self-attraction (i.e., compaction), protein-particle size, and protein concentration, we show that LADs behave as size-dependent steric environments: larger proteins are strongly hindered from entering dense peripheral chromatin, whereas smaller proteins retain limited access through nonspecific interactions. Notably, protein particles can interact with active chromatin regions at the boundary separating LADs from bulk. Overall, our simulations suggest a generic physical contribution to reduced regulatory access at the nuclear periphery while providing a biophysical mechanism for the transcriptional escape at largely gene-passive chromatin regions.

## II. METHODS

### A. Coarse-grained model of the nuclear periphery

To capture the essential physical features of the nuclear periphery environment, we constructed a coarse-grained model of a local, random patch of the nuclear periphery containing three components: the nuclear lamina, a chromatin polymer, and freely diffusing proteins. All components were represented by single or bonded spherical beads.

The characteristic length scales of the chromatin fiber (of diameter ∼10 nm, forming loops of size ∼ 10^2^ nm) and LAD thickness (∼10^2^ nm) are much smaller than the nuclear radius of curvature (∼10^3^ nm) [28–32]. For that reason, the lamina was approximated as a planar, static, impenetrable surface, and thus modeled as a fixed layer of beads at the bottom boundary of the simulation box at *z* = 0 (see Fig. 1).

**FIG. 1.**
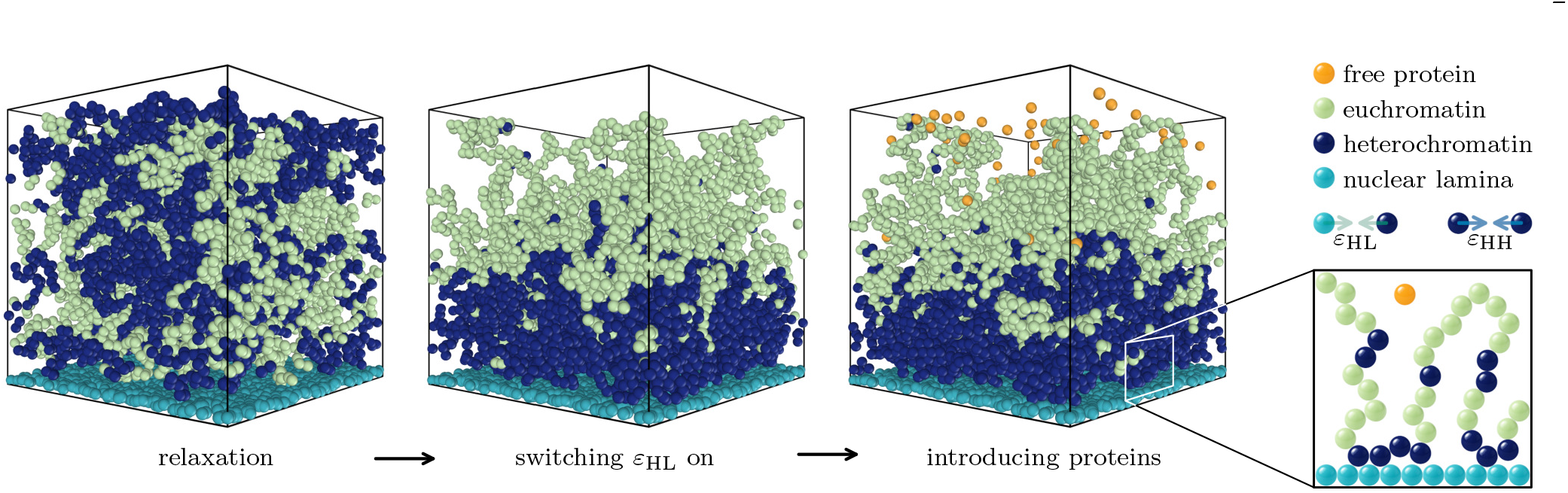
Simulation model and protocol. The nuclear lamina is represented as a fixed planar layer of beads at *z* = 0, chromatin is modeled as a bead–spring copolymer, and proteins are represented as single beads. The workflow consists of four stages: initial chromatin relaxation, activation of heterochromatin–lamina attraction, and protein insertion, before the production runs.

Diffusing proteins are modeled as single beads with various radii diffusing in the nucleoplasmic volume under Langevin dynamics. To isolate nonspecific steric effects, all protein–protein, protein–chromatin and protein–lamina interactions at a distance *r* are taken to be purely repulsive and described by a soft potential,

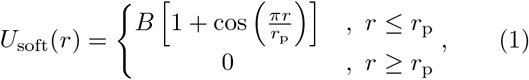

with interaction energy scale *B* = 20 *k*_B_*T* (where *k*_B_ is the Boltzmann constant and *T* the absolute temperature), and cut-off distance *r*_p_. By varying *r*_p_, we control the effective steric size of the diffusing proteins.

The chromatin chain is modeled using the Kremer–Grest bead–spring copolymer model [33]. The copolymer is composed of heterochromatin and euchromatin segments assigned using experimentally determined LAD maps based on ChIP-seq data [34] (see Fig. 1). This particular sequence corresponds to a coarse-grained version of human chromosome 20 with *N* = 6303 beads, with each bead corresponding to approximately 10 kb of DNA, or roughly 50 nucleosomes, with a size on the order of 10 nm.

Adjacent chromatin beads are bonded by the finitely extensible nonlinear elastic (FENE) potential,

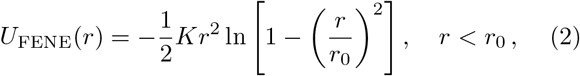

with bond stiffness *K* = 30 *k*_B_*T/σ*^2^ and maximum bond extension *r*_0_ = 1.5 *σ*. We employ *σ* as the basic simulation length unit. No explicit bending potential was included, so the chromatin chain is fully flexible at the coarse-grained level.

Nonbonded interactions between chromatin–chromatin and chromatin–lamina bead pairs are modeled using truncated-and-shifted Lennard–Jones (LJ) potential:

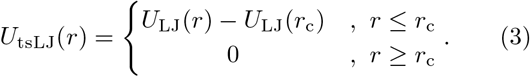

Here, *U*_LJ_(*r*) = 4*ϵ* (*r/σ*)^*−*12^ *−* (*r/σ*)^*−*6^ is the bare LJ potential, and *r*_c_ is the cutoff distance that we set to *r*_c_ = 2.5 *σ* in all simulations (except for pure repulsive cases, see below). We considered a range of interaction strengths between heterochromatin– lamina (*ϵ* = *ε*_HL_) and heterochromatin–heterochromatin (*ϵ* = *ε*_HH_) bead pairs to control the heterochromatin localization and compaction near the nuclear lamina. For euchromatin–heterochromatin, euchromatin– euchromatin and euchromatin–lamina interactions, we set *ϵ* = 1 *k*_B_*T* . Purely repulsive interactions are implemented using the Weeks–Chandler–Andersen cutoff *r*_*c*_ = 2^1*/*6^*σ* with *ϵ* = 1 *k*_B_*T* .

### B. Molecular dynamics simulation protocol

All molecular dynamics simulations are performed with lammps [35] in the canonical ensemble (constant *NV T* ) using a Langevin thermostat. Data analysis used Pandas [36], and visualization was carried out with ovito [37]. The integration time step is Δ*t* = 0.0005 *τ*, where 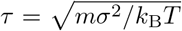 is the simulation time scale and *m* is the mass for all types of simulated beads. The simulation workflow consist of four stages as illustrated in Fig. 1.

First, the initial random chromatin configuration is relaxed for 2 ×10^6^ integration steps in a simulation box spanning 0 ≤ *z* ≤ 35 *σ* in the direction normal to the lamina and −17.5≤ *x, y* ≤17.5 *σ* in lateral directions. During this stage, all nonbonded interactions are purely repulsive.

Second, heterochromatin–lamina attraction *ε*_HL_ is switched on, and the system was evolved for 2 ×10^7^ steps to allow chromatin adsorption and reorganization at the lamina.

Third, proteins are inserted into the simulation box. To reduce overlap, the upper boundary of the simulation box is temporarily shifted from *z* = 35 *σ* to *z* = 68 *σ*, and proteins are randomly placed in a region spanning 33.5 *σ*≤ *z* ≤ 62.5 *σ* in the direction normal to the lamina and −12.5 *σ* ≤ *x, y* 12.5 ≤ *σ* in lateral directions. The system is then evolved for 1 ×10^5^ steps using soft interactions (*U*_soft_) for all bead pairs to remove overlaps, followed by another 1×10^5^ steps with purely repulsive interactions to redistribute the proteins. At the end of this stage, the upper boundary of the simulation box is returned to its original position over 2×10^5^ steps, after which, protein-involved interactions were restored to *U*_soft_ and the remaining interactions are set back to the target truncated-shifted Lennard-Jones (*U*_tsLJ_) forms. To generate different chromatin environ-ments, various cases of heterochromatin–lamina interactions (*ε*_HL_) and heterochromatin–heterochromatin self-interactions (*ε*_HH_) are explored, see Table I.

**TABLE 1.** Ranges scanned for the heterochromatin self-interaction *ε*_HH_, heterochromatin–lamina attraction *ε*_HL_, protein size *r*_P_, and overall protein density *c*_P_.

| Parameter | Range |
| --- | --- |
| $\epsilon_{\text{HH}}$ | $[0.10, 1.10] k_B T$ & repulsive |
| $\epsilon_{\text{HL}}$ | $[0.35, 1.40] k_B T$ |
| $r_{\text{P}}$ | $[0.25, 4.00] \sigma$ |
| $c_{\text{P}}$ | $[0.00117, 0.01866] \sigma^{-3}$ |

Finally, 10 independent replicas with different random-number seeds were run for each parameter set. Production runs were carried out for 2 × 10^6^ steps, and time-averaged density profiles were computed over the last 1 ×10^6^ steps. At the end, densities were averaged over replicas, and the error bars were obtained via the standard deviation across replicas (see, *e*.*g*., Fig. 2).

**FIG. 2.**
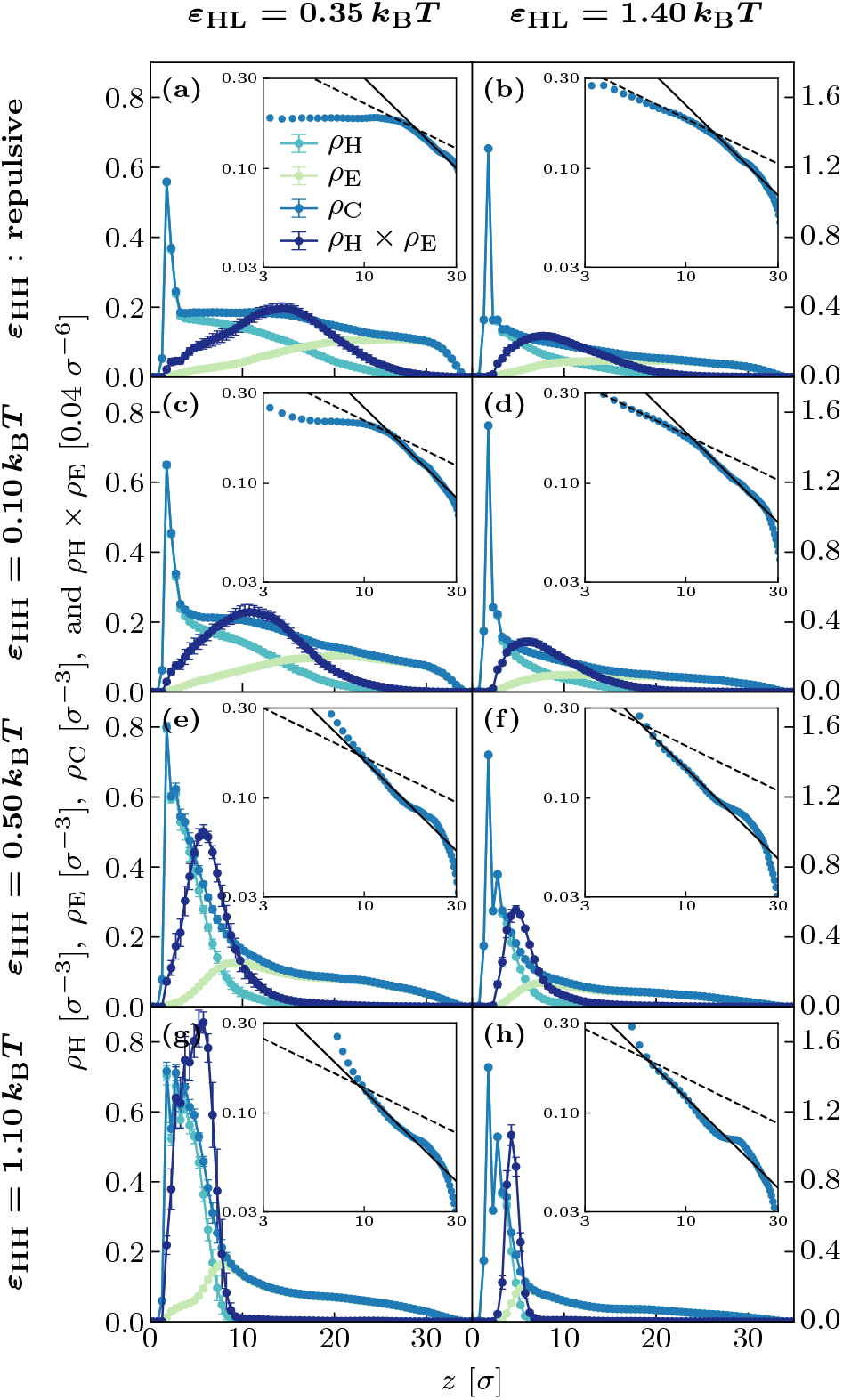
Heterochromatin, euchromatin, and total chromatin density profiles, *ρ*_H_(*z*), *ρ*_E_(*z*), and *ρ*_C_(*z*), together with the density-overlap profile, *ρ*_H_(*z*) × *ρ*_E_(*z*), in the absence of proteins. From top to bottom, the rows correspond to repulsive, weakly attractive, moderately attractive, and more strongly attractive heterochromatin–heterochromatin interactions (*ε*_HH_); while the left and right columns respectively correspond to weaker and stronger heterochromatin–lamina attractions (*ε*_HL_), see the row and column headings. Insets show *ρ*_C_(*z*) on log–log scales with *z*^*−*1*/*2^ and *z*^*−*1^ guide lines.

### C. Definition of the LAD boundary

To determine the spatial extent of the LADs, one-dimensional number-density profiles *ρ*_H_(*z*) of heterochromatin and *ρ*_E_(*z*) of euchromatin along the direction normal to the lamina is computed. The simulation box was divided into 70 bins, each of height 0.5 *σ* along the *z* axis, and bead densities were calculated and time-averaged for each bin (see Fig. 2).

A density-overlap profile using the Hadamard product *ρ*_H_(*z*) ×*ρ*_E_(*z*) of the heterochromatin and euchromatin density profiles is determined. Figure 2 shows this product for representative values of *ε*_HH_ and *ε*_HL_ in the absence of proteins. This density-overlap is small in regions dominated by only one chromatin type and becomes larger where both types coexist appreciably. We therefore used the peak position *z* = *L*^lad^ of the product *ρ*_H_(*z*) ×*ρ*_E_(*z*) as an operational, datadriven definition of the boundary between the lamina-associated heterochromatin-rich region and the interior euchromatin-rich region. Bins between the lamina and this peak position (0≤ *z* ≤*L*^Lad^) were classified as belonging to the LAD region.

After locating the position of the LAD boundary, chromatin densities and protein occupancies are integrated from the lamina surface up to this boundary for each protein size, overall protein concentration, and interaction parameter set (see Table I). These integrated quantities are used to quantify protein penetration into LADs and to compare steric accessibility across different chromatin environments.

## III. RESULTS

### A. Chromatin organization in the absence of proteins

Various tethering proteins can increase the local density of chromatin near the nuclear periphery [38–42]. To this extent, before introducing proteins into the system, we first characterize the dependence of chromatin density near the periphery on heterochromatin–lamina attraction (*ε*_HL_) and also heterochromatin self-interaction (*ε*_HH_). Figure 2 shows representative one-dimensional density profiles of heterochromatin *ρ*_H_(*z*), euchromatin *ρ*_E_(*z*), and total chromatin *ρ*_C_(*z*) for different combinations of *ε*_HH_ and *ε*_HL_, where *z* is the distance fron the surface. In all cases, heterochromatin is preferentially localized near the lamina due to attractive *ε*_HL_, while euchromatin is displaced toward larger distances from the lamina. Increasing heterochromatin self-attraction strongly reshapes the partitioning of chromatin species near the lamina. As *ε*_HH_ is increased, heterochromatin becomes more compact and accumulates more sharply within the peripheral LAD. Consequently, the overlap region between heterochromatin-rich and euchromatin-rich domains narrows, and the interface between these regions becomes more pronounced. These trends are enhanced at larger heterochromatin–lamina affinity, *ε*_HL_, indicating that heterochromatin self-attraction and lamina binding cooperate in setting the structure and sharpness of the LAD environment.

The inset plots in Fig. 2 show that, over the range of parameters considered here, the total chromatin density profile follows a power-law decay away from the lamina. In all cases of *ε*_HH_ and *ε*_HL_, the total chromatin density is consistent with the power-law form *ρ*_C_(*z*) ∼ *z*^*−*1^ (see insets in Fig. 2), as predicted by scaling theory for adsorption of *linear* polymers [43] and observed experimentally for, e.g., human T-cells [41]. Although a *ρ*_C_(*z*) ∼*z*^*−*1*/*2^ decay has also been predicted by scaling theory for adsorption of *cyclic* polymers and reported experimentally for several eukaryotic cell types [43], in the present simulations this behavior is limited to narrower portions of selected profiles, whereas the −1 decay provides a consistent description across the cases shown in Fig. 2.

Across all interaction strengths, the total chromatin density reflects the combined effects of lamina affinity and heterochromatin self-interaction, showing a progressive transition from a diffuse chromatin organization to a strongly layered structure as *ε*_HH_ and *ε*_HL_ are increased. These results, in the absence of proteins, demonstrate that heterochromatin self-attraction and lamina binding act cooperatively to regulate the spatial extent and sharpness of LADs, the chromatin density inside LADs, and hence the chromatin mesh size (porosity) and available steric volume to other molecules inside LADs.

To quantify these trends, we next determine the LAD thickness *L*^lad^ and compute the average total chromatin density 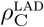 inside the LAD for various values of *ε*_HH_ and *ε*_HL_ (see Fig. 3). The LAD thickness decreases while the total chromatin density in the LAD increases systematically as either *ε*_HH_ or *ε*_HL_ is strengthened (see Fig. 3), showing that stronger heterochromatin self-attraction and stronger lamina binding both reduce the sterically accessible volume within LADs. In other words, *ε*_HH_ and *ε*_HL_ do not merely shift the position of the peripheral chromatin layer; they also control its local packing density and therefore its effective porosity.

**FIG. 3.**
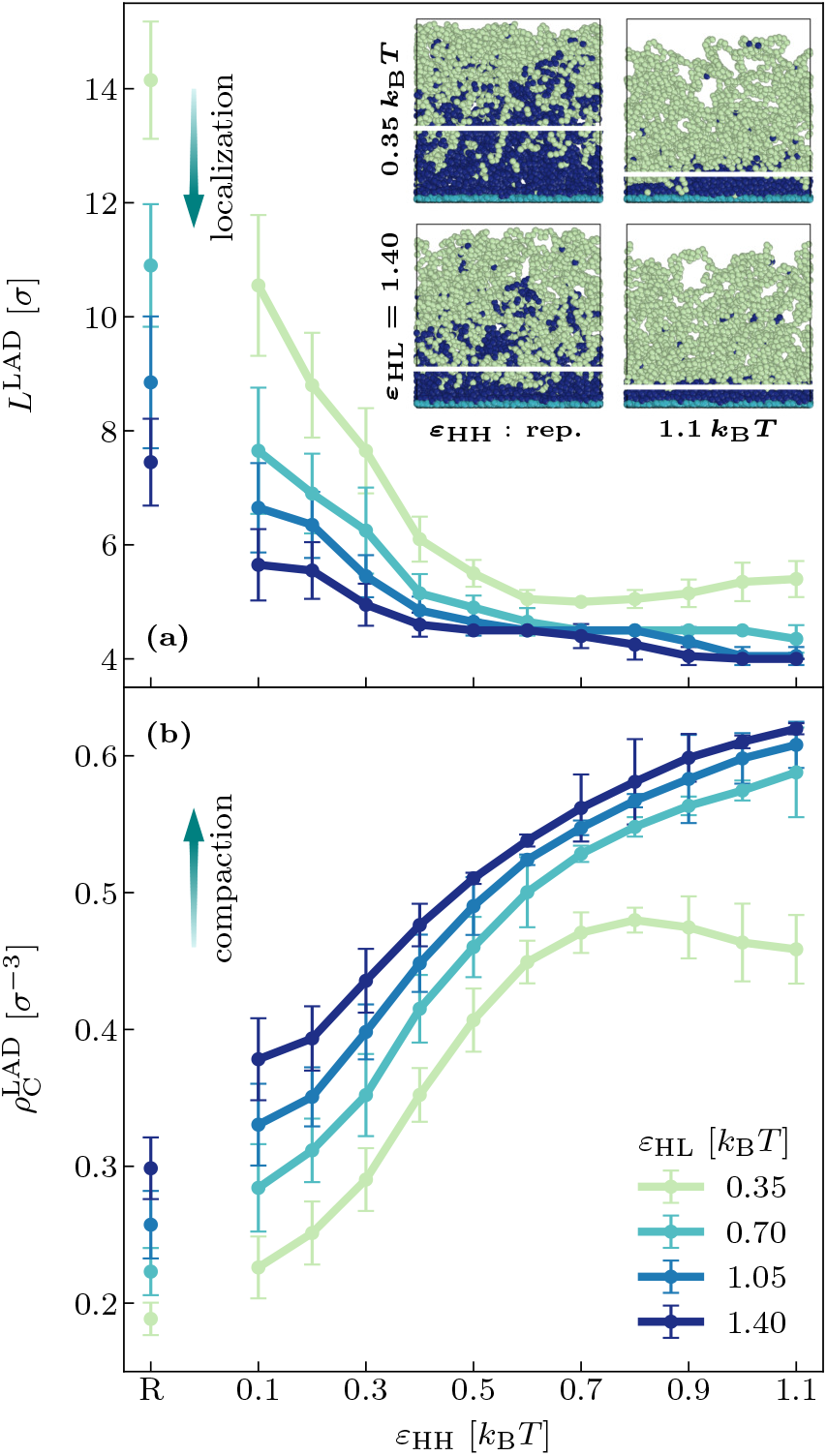
**(a)** LAD thickness *L*^lad^, and **(b)** total chromatin density inside LAD 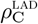 as functions of *ε*_HH_ for different values of *ε*_HL_. Results for repulsive *ε*_HH_ are also shown as separate points as indicated by “R” on the horizontal axis. Insets show representative simulation snapshots for interaction pairs at the extremes of the parameter ranges explored: repulsive and strongly attractive *ε*_HH_ in left and right inset panels, and weakly and strongly attractive *ε*_HL_ in top and bottom inset panels. White horizontal lines mark average LAD boundaries.

Taken together, these results show that the chromatin environment at LADs is governed by two coupled physical controls: heterochromatin self-attraction, which promotes compaction, and heterochromatin-lamina attraction, which favors peripheral localization [44]. The observables *L*^Lad^ and 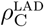 therefore provide a compact structural description of the background against which protein partitioning should be interpreted.

### B. Chromatin organization is insensitive to protein presence, size, and concentration

Next, we characterize how added protein particles can influence the LAD structure. To this end, we run separate simualtions and determine the LAD thickness (*L*^lad^) and the total chromatin density within the LAD 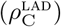 in the presence of proteins of different sizes *r*_P_ and at various overall protein concentrations *c*_P_. We observed that the chromatin environment established by *ε*_HH_ and *ε*_HL_ is not measurably altered by the proteins introduced in our simulations, with changes being within the statistical error margins.

Thus, in the parameter range explored in this work (see Table I), proteins do not reorganize or swell the chromatin background and do not modify the spatial extent or composition of the LAD region significantly. Instead, as discussed in the previous subsection, the chromatin structure is set primarily by the heterochromatin self-interaction *ε*_HH_ and the heterochromatin–lamina attraction *ε*_HL_. This interpretation is consistent with the simulation protocol design, in which different chromatin environments are generated first and protein insertion is performed afterward.

Since the chromatin observables are insensitive to the presence, size, and overall concentration of the protein particles, modeled as soft steric spheres in our study, the proteins act as passive probes of a pre-existing LAD environment rather than as agents that reshape it. The dependence of protein occupancy on their size *r*_P_ and overall concentration *c*_P_ can therefore be interpreted directly in terms of steric partitioning within a fixed chromatin background. For this reason, in the following subsections, we focus on LAD occupancy of proteins, while treating the chromatin environment as externally controlled by *ε*_HH_ and *ε*_HL_ alone.

### C. Size-dependent steric partitioning of proteins into LADs

Having established that proteins do not alter the chromatin background in a statistically significant manner in our calculations, we next examine how the pre-existing LAD environment affects protein partitioning as the protein size *r*_P_ is varied at fixed overall protein concentration *c*_P_.

This size dependence of protein exclusion is quantified in Fig. 4 through the average protein density within the LAD region, 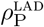. For every chromatin environment considered, 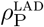 decreases with increasing *r*_P_, confirming that LADs act as size-selective steric barriers. Moreover, the extent of depletion depends on the chromatin environment itself: denser and more strongly lamina-localized LADs produce stronger suppression of protein occupancy. Thus, the effect of protein partitioning at a fixed overall concentration is controlled jointly by the protein size *r*_P_ and by the chromatin density and organization set by *ε*_HH_ and *ε*_HL_.

**FIG. 4.**
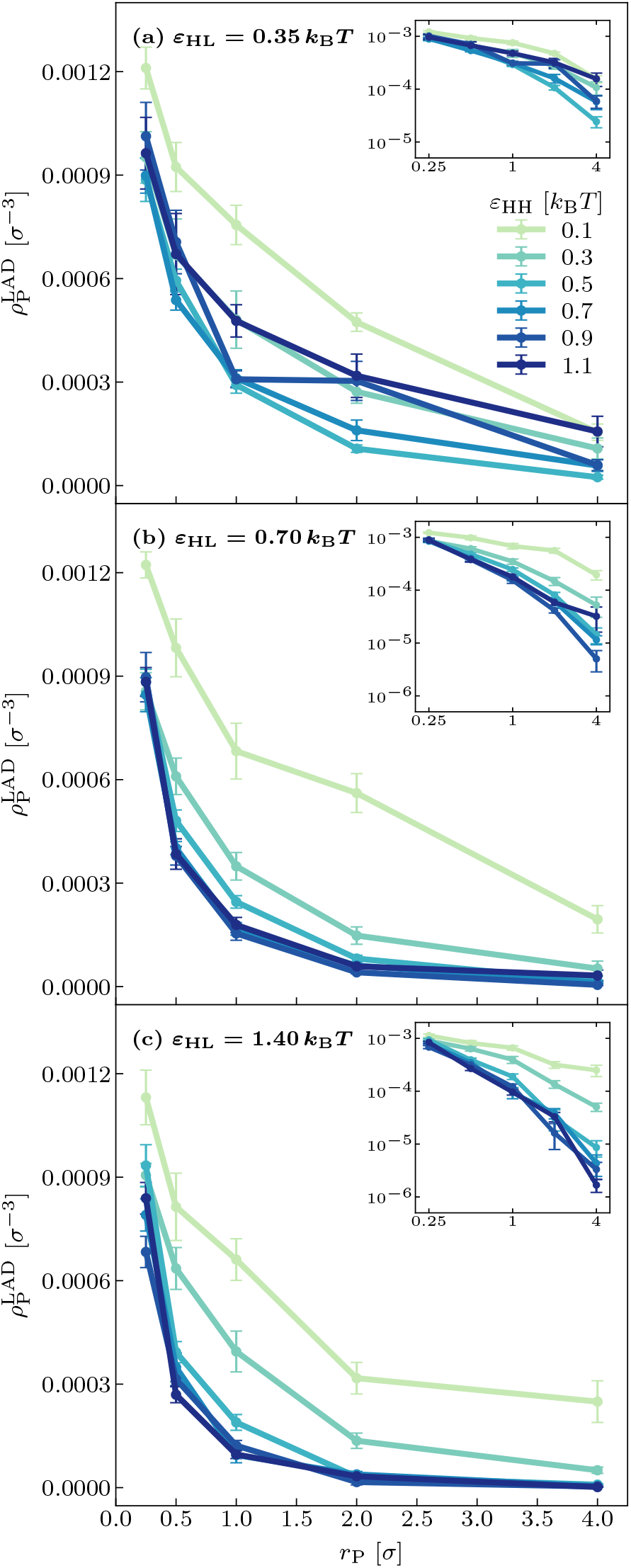
Protein densities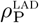 inside LADs as functions of protein size *r*_P_ plotted for various *ε*_HH_for a represantative overall protein concentration *c*_P_ = 0.00117 *σ*^*−*3^, and **(a)** *ε*_HL_ = 0.35 *k*_B_*T*, **(b)** *ε*_HL_ = 0.70 *k*_B_*T*, **(c)** *ε*_HL_ = 1.40 *k*_B_*T* . Insets show the data in log–log axes. *titioning environments*, whose filtering strength depends on the underlying chromatin compaction.

Taken together, these results show that the steric effect of LADs is graded rather than binary. Small proteins are not completely free of hindrance, but they retain limited access to the peripheral chromatin layer, whereas larger proteins are increasingly excluded as their effective size increases. Therefore, proteins are not merely hindered by LADs, but LADs behave as *size-selective steric partitioning environments*, whose filtering strength depends on the underlying chromatin compaction.

### D. Protein density profiles collapse when normalized by the overall protein concentration

We next examine how the protein distributions depend on the overall protein concentration *c*_P_ at fixed protein size *r*_P_. Figure 5(a) shows that increasing *c*_P_ increases the protein density profile *ρ*_P_(*z*) essentially uniformly throughout the system. The overall amplitude of the profile grows with *c*_P_, but its shape remains unchanged. In particular, the depletion of proteins near the lamina and the suppression of protein density within the LAD region persist over the whole concentration range considered here. Thus, changing *c*_P_ does not qualitatively alter the spatial pattern of protein partitioning generated by the local chromatin environment.

**FIG. 5.**
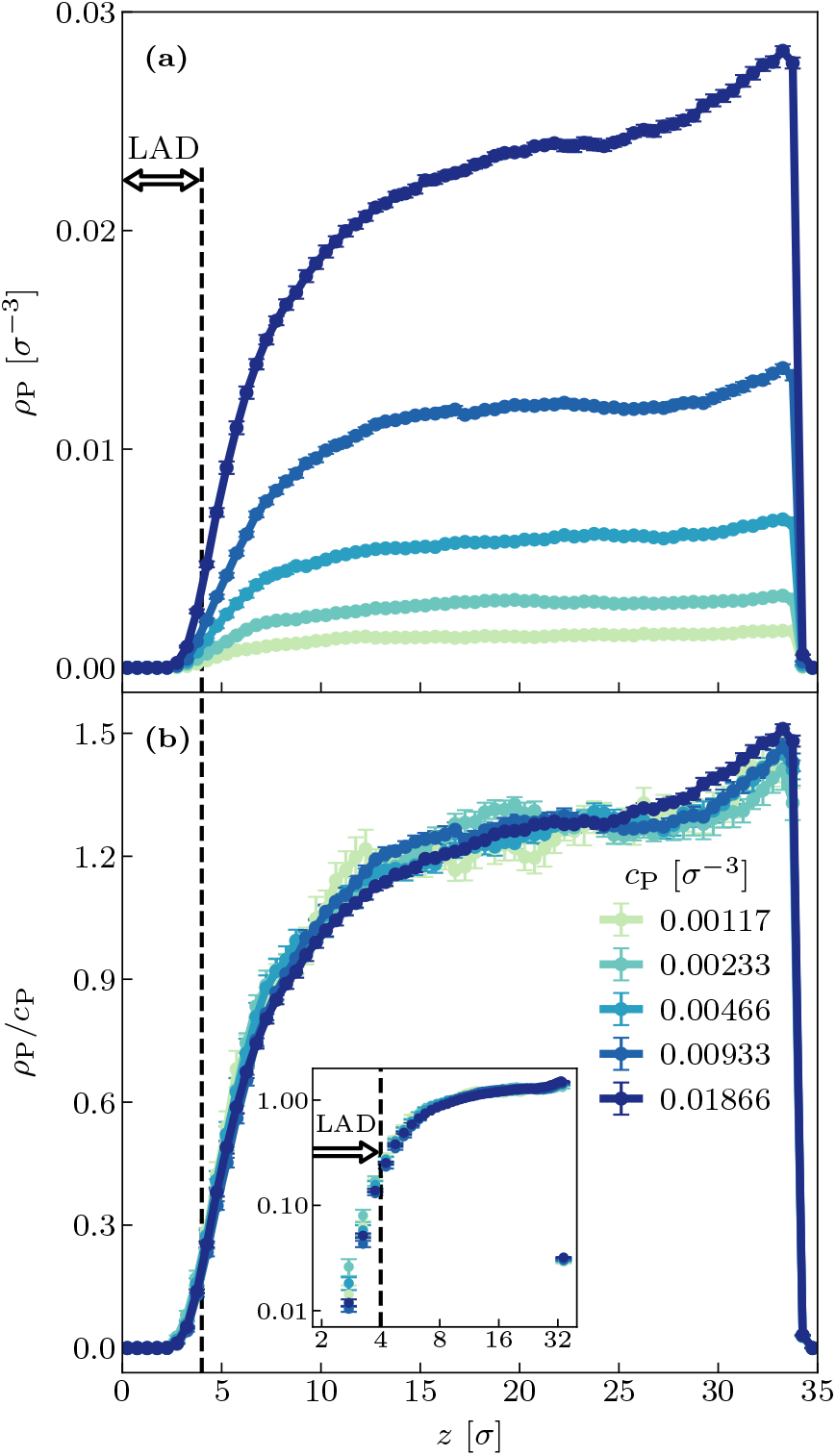
**(a)** One-dimensional protein density profiles, *ρ*_P_(*z*), for different overall protein concentrations *c*_P_, at fixed protein size *r*_P_ = 1 *σ* and fixed chromatin interaction parameters *ε*_HH_ = 1.1 *k*_B_*T* and *ε*_HL_ = 1.4 *k*_B_*T* . Increasing *c*_P_ increases the overall amplitude of the protein profile without substantially changing its shape. **(b)** The rescaled profiles *ρ*_P_(*z*)*/c*_P_ collapse onto a master curve, showing that protein partitioning is linear in concentration and that the spatial profile is controlled primarily by the chromatin environment and protein size. The inset panel shows the data in log-log axes. Dashed vertical lines in all panels mark the LAD boundary.

This becomes more evident when the protein density profiles are normalized by the overall protein concentration *c*_P_. As shown in Fig. 5(b), the rescaled profiles *ρ*_P_(*z*)*/c*_P_ collapse onto a master curve for all values of *c*_P_. The collapse indicates that, within the regime studied here, the protein distribution is linear in concentration: the chromatin environment and the protein size determine the shape of the profile, while *c*_P_ sets only its overall magnitude. Equivalently, the normalized protein occupancy inside the LAD region,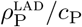, is indepen-dent of *c*_P_ within numerical uncertainty.

This finding shows that the concentration dependence of protein partitioning is essentially trivial in the present model and that the nontrivial control parameters are instead the protein size *r*_P_ and the chromatin environment generated by *ε*_HH_ and *ε*_HL_. In other words, the steric filtering effect of LADs is already encoded in the chromatin structure, and varying the total number of proteins mainly rescales the population of particles exploring that same steric landscape.

### E. Scaling collapse of LAD protein partitioning

Solute partitioning in polymeric media depends on solute size, polymer architecture, density, and mesh size [45]. Our results show that the normalized protein occupancy within LADs is controlled by protein size and the chromatin environment, whereas the overall protein concentration only rescales the density profiles. This suggests that protein penetration into LADs can be described as an equilibrium partitioning problem. Hence, we define the protein–LAD partitioning coefficient, as the Boltzmann weight

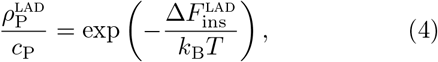

where 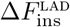is the free energy cost of inserting a finite-size protein particle into the LAD. In polymeric media, this free energy cost can generally contain osmotic, interfacial, and elastic contributions. The osmotic contribution is proportional to the work needed to create a cavity of volume 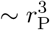 within LADs, while the interfacial contribution is proportional to the particle surface area 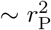 . Thus, at the scaling level,

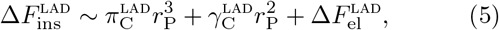

where 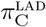 and 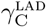 are the osmotic pressure and surface tension of the LAD chromatin environment, respectively.

Such volumetric and areal terms are standard in cavity-insertion descriptions of particles in polymer brushes [46], polymer networks [47], and dilute or semidilute polymer solutions [48].

In the present simulations, however, the protein particles penetrate a concentrated chromatin layer adsorbed to the nuclear lamina. We therefore assume that the dominant contribution over the explored parameter range is an elastic-mesh penalty associated with deforming the transient steric constraints of the LAD chromatin network. We write this contribution in the scaling form

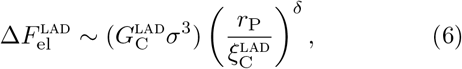

where 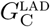 is an effective elastic constraint modulus, 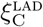is the LAD mesh size, and *δ* is an exponent reflecting the elasticity of chromatin steric constraints. For the LAD layer in concentrated regime, we estimate the effective constraint modulus as proportional to the density of chromatin constraints, 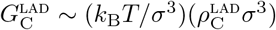, and the mesh size as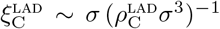[49]. With these estimates, the elastic-mesh insertion free energy becomes 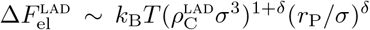, and the LAD partition coefficient is predicted to obey

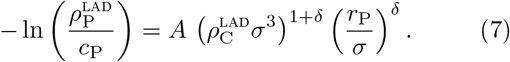

Figure 6 shows the resulting scaling collapse of our simualtion data obtained for different heterochromatin self-attractions *ε*_HH_, heterochromatin-lamina attractions *ε*_HL_, and protein sizes *r*_P_ (see Fig. 4). The data collapse is obtained for the fitted prefactor *A* = 5.254 ±0.154, and the fitted exponent

**FIG. 6.**
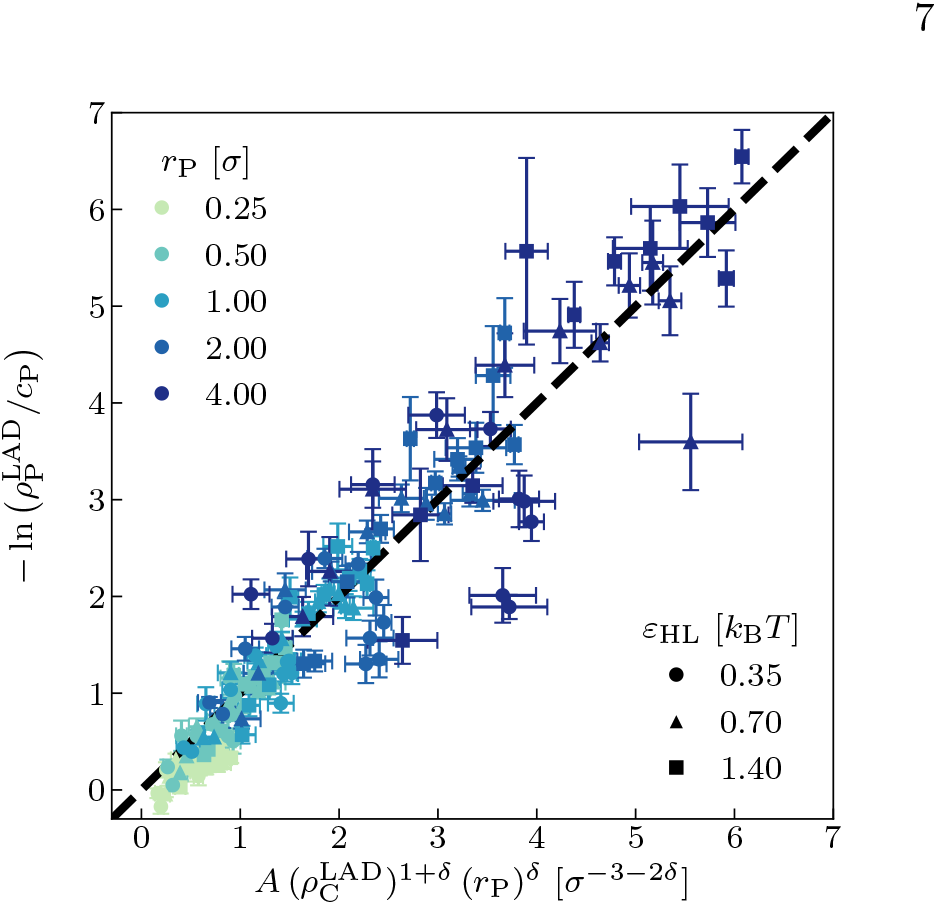
The dimensionless LAD partitioning free energy, 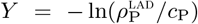, plotted against the elastic-mesh scaling variable 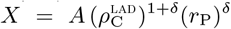, see Eq. 7, for overall protein concentration *c*_P_ = 0.00117 *σ*^*−*3^. Different *r*_P_ are coded with different colors while different *ε*_HL_ are coded with different symbols. Each data set includes 11 points for *ε*_HH_ = 0.1, 0.2, …, 1.1 *k*_B_*T* . Data from different chromatin interaction parameters and protein sizes collapse onto the dashed universal line *Y* = *X* for fitted parameters *A* = 5.254 and *δ* = 0.687.

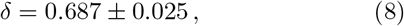

where the errors indicate the standard deviations of the fit process. Interestingly, the fitted exponent is very close to exponents reported in other polymeric sieving environments. In partitioning of ribonucleoprotein complexes from the cellular actin cortex, an excluded-volume model of the actin cortex meshwork explains the observed depletion, with a fitted particle size exponent [50]. A similar numerical value was also reported for electrophoretic retardation of near-spherical proteins in semidilute polyethylene glycol solutions, where a concentration exponent 0.69 was found [51]. Although these systems differ from the LAD chromatin, the recurring value ≈0.7 suggests that sublinear steric penalties may be a common feature of macromolecular penetration into polymeric media. This sublinear exponent indicates that the protein-insertion penalty is not governed simply by the bare particle volume or surface area. Instead, protein exclusion from LADs is controlled by the transient polymeric mesh of the chromatin layer. Increasing *ε*_HH_ or *ε*_HL_ increases the density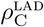, reduces the mesh size 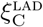, and thereby raises the free-energy cost of protein penetration, which also increases with the protein size *r*_P_. Consequently, the steric filtering effect of LADs can be expressed as an elastic-mesh partitioning law in which chromatin compaction and protein size enter through a single scaling variable.

## IV. DISCUSSION

The chromatin confined in the cell nucleus interacts with a physical interface, the nuclear envelope. This interface not only provides an impermeable border for the genome but also regulates and organizes the chromatin environment. Our simulations revealed that lamina-associated domains (LADs) may behave as size-selective steric filters that hinder the access of larger-sized proteins to the LADs, resulting from their association with the nuclear surface and self-compaction of chromatin. This filter only allows the entrance of relatively small proteins. As the chromatin environment at LADs becomes more compact and dense, fewer proteins can permeate LADs and approach the nuclear surface. As a result, the compaction of the chromatin environment affects the hindrance effect felt by the transcription proteins. The mechanism of this hindrance is nonspecific, as the only interaction between the proteins and LADs considered in our modeling is a nonspecific, pairwise interaction potential, see Eq. (1).

Experimentally, RNA Polymerase II, which is a large protein complex of size 10 –15 nm, has been shown to interact with chromatin near the nuclear periphery minimally [11, 52]. Our results align with this experimental observation, suggesting that such large proteins may be excluded from LADs due to the steric hindrance mechanism demonstrated here. Relatedly, synthetic dextran particles of sizes around 5 –10 nm were shown to proximite the nuclear periphery, whereas larger particles were excluded from the nuclear boundary [53]. In addition, a recent experimental study indicates that both euchromatin and heterochromatin are highly accessible in living human cells and transcriptional regulation may rely more on transcription protein presence [27]. Our results support this key finding and suggest that the lower expression levels in the LADs could be correlated with the lack of transcription proteins inside the LADs.

Importantly, our results show that at the boundary of LADs, the protein concentration does not vanish abruptly but rather follows an S curve pattern (see Fig. 5). This suggests that there is a finite number of proteins that can interact with LADs and possibly with open chromatin segments (if any) buried inside LADs. Notably, even though larger proteins have limited access to LADs, small transcription factors (TF) can enter this layer and bind to DNA. Diffusing of such TF-DNA complexes towards the LAD border may allow Polymerase binding to these complexes, resulting in a transcriptional “escape” mechanism [23, 24].

While our coarse-grained molecular dynamics approach captures the fundamental-physics aspects of transcriptional proteins’ escape into LADs, it does not account for specific interactions between various types of proteins and ignores histone modifications. Also, the model only captures the dynamics for a small section of the nuclear periphery environment, and it does not include the dynamics of lamina proteins, as the model assumes the lamina as a static layer. In reality, lamin protein dynamics may affect the chromatin organization near the periphery [15]. Although our simulations contribute insights into transcription suppression, we ignore the active processes driven by motor-protein activity. Incorporating these factors in future simulations could clarify how LADs’ mobility couples to gene regulation.

Overall, our study provides a polymer-physics-based understanding of the role of LADs in transcription. Our results show that one key role of LADs could be the steric hindrance of transcription factors by their polymeric nature, which can regulate gene inactivity in heterochromatin-rich LADs. These results may stimulate future studies and provide a basis for the development of large-scale, more detailed models incorporating lamin dynamics and enzymatic activity.

## ACKNOWLEDGMENTS

This work was supported by TÜ BİTAK, The Scientific and Technological Research Council of Türkiye (Grant no. 122F309), and by Scientific Research Projects Commission of Pîrî Reis University (Project no. BAP-2024-02-02). The numerical calculations reported in this work were partially performed at the Tü BİTAK ULAKBİM High Performance and Grid Computing Center (TRUBA Resources).

## Notes

https://zenodo.org/records/19662033?token=eyJhbGciOiJIUzUxMiJ9.eyJpZCI6IjQ1MDI0ZGYxLTA4YjItNDQxNS05MDU3LTFhNmIyYzMxNjEyZSIsImRhdGEiOnt9LCJyYW5kb20iOiJkNTVmN2ZhNDhlZTY1ZmJlOWU0MjUzN2I0ZDgxOTNhZiJ9.qWTD0cPZrlgzMPLTcljKmngbt_kR82GuEc0kmbGCGj-1pJR2S_qpSitJE1uzH8NGetR9QXzbB_99hTrwe6gzjA

### Competing Interest Statement

The authors have declared no competing interest.

